# Single-particle cryo-EM at atomic resolution

**DOI:** 10.1101/2020.05.22.110189

**Authors:** Takanori Nakane, Abhay Kotecha, Andrija Sente, Greg McMullan, Simonas Masiulis, Patricia M.G.E. Brown, Ioana T. Grigoras, Lina Malinauskaite, Tomas Malinauskas, Jonas Miehling, Lingbo Yu, Dimple Karia, Evgeniya V. Pechnikova, Erwin de Jong, Jeroen Keizer, Maarten Bischoff, Jamie McCormack, Peter Tiemeijer, Steven W. Hardwick, Dimitri Y. Chirgadze, Garib Murshudov, A. Radu Aricescu, Sjors H.W. Scheres

## Abstract

The three-dimensional positions of atoms in protein molecules define their structure and provide mechanistic insights into the roles they perform in complex biological processes. The more precisely atomic coordinates are determined, the more chemical information can be derived and the more knowledge about protein function may be inferred. With breakthroughs in electron detection and image processing technology, electron cryo-microscopy (cryo-EM) single-particle analysis has yielded protein structures with increasing levels of detail in recent years^1,2^. However, obtaining cryo-EM reconstructions with sufficient resolution to visualise individual atoms in proteins has thus far been elusive. Here, we show that using a new electron source, energy filter and camera, we obtained a 1.7 Å resolution cryo-EM reconstruction for a prototypical human membrane protein, the β3 GABA_A_ receptor homopentamer3. Such maps allow a detailed understanding of small molecule coordination, visualisation of solvent molecules and alternative conformations for multiple amino acids, as well as unambiguous building of ordered acidic side chains and glycans. Applied to mouse apo-ferritin, our strategy led to a 1.2 Å resolution reconstruction that, for the first time, offers a genuine atomic resolution view of a protein molecule using single particle cryo-EM. Moreover, the scattering potential from many hydrogen atoms can be visualised in difference maps, allowing a direct analysis of hydrogen bonding networks. Combination of the technological advances described here with further approaches to accelerate data acquisition and improve sample quality provide a route towards routine application of cryo-EM in high-throughput screening of small molecule modulators and structure-based drug discovery.

## Introduction

Multiple factors determine the attainable resolution of reconstructions from cryo-EM images of individual biological macromolecules and their complexes^4^. However, the radiation damage caused by electron interactions with the sample is a fundamental limitation. In order to preserve the molecular structure, damage is restricted by carefully limiting the number of electrons used for imaging^5^. The resulting counting statistics lead to high levels of experimental noise. Moreover, the biological molecules are suspended in a thin layer of vitrified water, which contributes additional noise in the images. The signal-to-noise ratio (SNR) of cryo-EM images drops rapidly with spatial frequency, and at higher spatial frequencies the noise is typically orders of magnitude higher than the signal.

High-resolution reconstructions may still be calculated by combining many cryo-EM images. DeRosier and Klug showed that the two-dimensional Fourier transform of each individual image, or particle, represents a central slice through the three-dimensional Fourier transform of the original object^6^. Therefore, provided their relative orientations can be determined, one can reduce the amount of noise in the reconstruction by averaging over multiple particles in Fourier space. However, because the reduction of noise in the average only scales with the square root of the number of observations, acquiring more particles is often less efficient than increasing the SNR of the individual particles. Increasing the SNR of the particles also improves the accuracy with which their orientations can be determined. Consequently, although microscope automation^7^ and faster image processing programs^8,9^ have allowed reconstructions from larger data sets in recent years, greater benefits may be achieved through increasing the SNR of the raw data^10^.

Around 2013, two technological developments were of particular importance in allowing a sudden increase in the resolution of cryo-EM reconstructions. Firstly, direct electron cameras with improved detective quantum efficiency compared to previously used photographic film led to a direct increase of SNR in cryo-EM images^11^. Secondly, fast electronics in these cameras provided functionality to record movies instead of the still images that were previously recorded on photographic film. This opened up the possibility to correct for movements in the vitrified sample that are caused by interactions with the electron beam, thereby further increasing SNRs^12–14^.

Here, we report the use of three new technological developments that further increase the SNR of cryo-EM images (**Figure 1**). Specifically, we used a new electron source, in the form of a cold field emission electron gun (FEG) that is optimised for energy spread and improves SNRs at resolutions better than 2.5 Å. In addition, we used the latest generation Falcon camera coupled to a new energy filter that removes inelastically scattered electrons. We demonstrate that these three developments lead to a marked increase in the achievable resolution that ultimately enables the visualisation of individual protein atoms in optimised samples.

**Figure 1:**
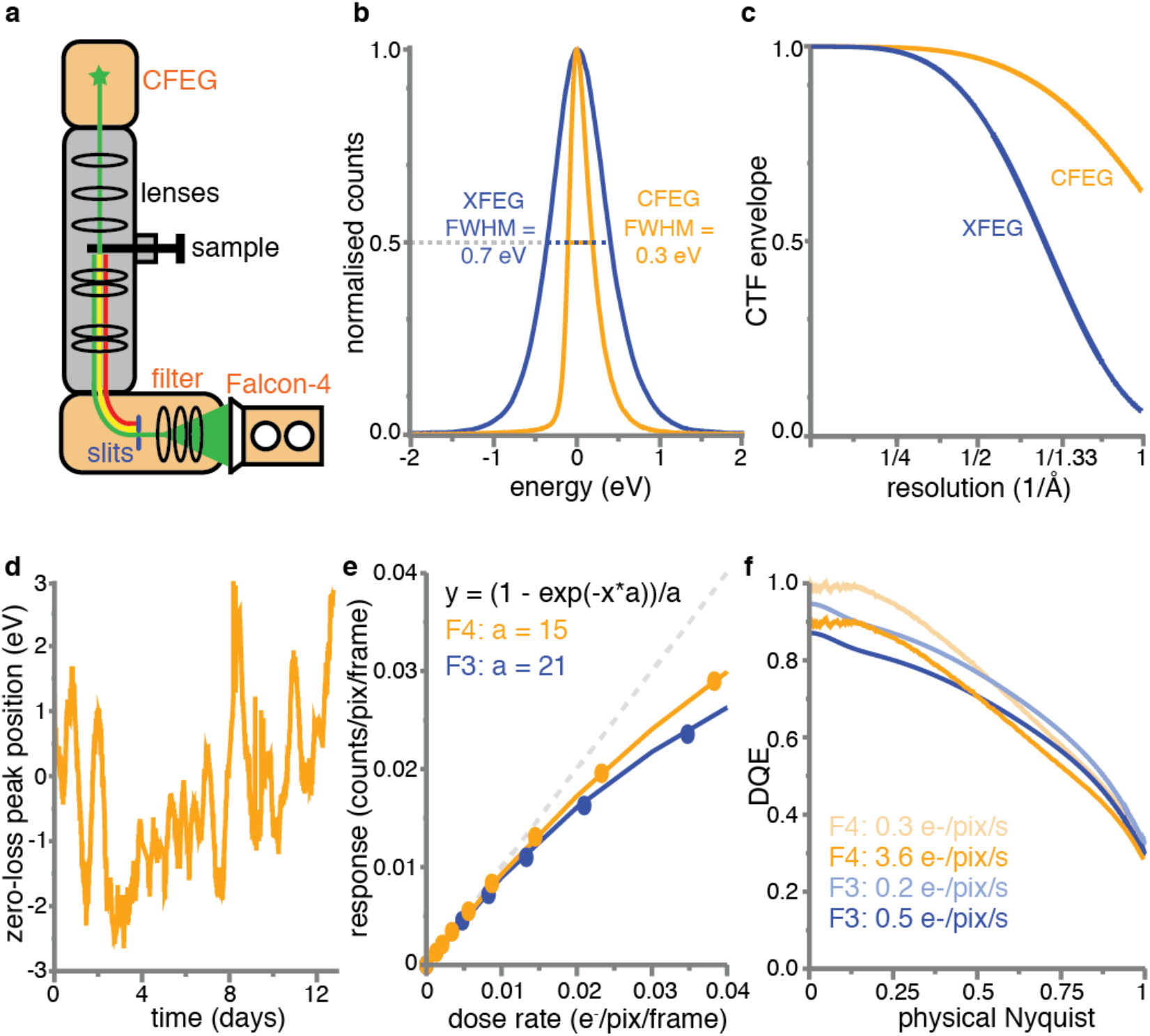
New imaging technologies for cryo-EM. **(a)** Schematic overview of an electron cryo-microscope. The new cold field-emission gun (CFEG), energy filter and Falcon-4 camera are highlighted in orange. **(b)** Energy spread of the XFEG (blue) and the CFEG (orange), with the full width of the curves at half their maximum value (FWHM). **(c)** Theoretical CTF envelope functions for the XFEG (blue) and the CFEG (orange). **(d)** The relative position of the zero-loss peak with respect to the centre of the slits in the energy filter over multiple days of operation. **(e)** Dose response measurements for the Falcon-3 (blue dots) and Falcon-4 (orange dots). The orange and blue lines are the corresponding fits to the data, with the fit parameters indicated in the same colours; the dashed grey line represents the perfect response. **(f)** DQE curves for the Falcon-3 at a dose rate of 0.2 (light blue) and 0.5 e^−^/pixel/s (blue) and the Falcon-4 at a dose rate of 0.3 (light orange) and 3.6 e^−^/pixel/s (orange).

### An electron source optimised for energy spread

The source inside the microscope generates electrons with a range of different wavelengths, or energies. Because not all these electrons can be focused in the same plane due to chromatic aberration in the objective lens, the energy spread of the electrons leads to a blur in the images. The corresponding loss in SNR increases with spatial frequency and is described by an envelope on the contrast transfer function (CTF) of the optical system. Many state-of-the-art electron microscopes are equipped with a field emission gun (FEG) that is operated at a temperature of 1700-1800 K and optimised for brightness (XFEG). In order to optimise the source for its energy spread, a so-called “cold” FEG (CFEG) may be operated at room temperature instead^15^. We used the prototype of a CFEG that was recently developed by Thermo Fisher Scientific (TFS) and is operated at a low beam current. Compared to the energy spread of 0.7 eV for a standard TFS XFEG, the new CFEG has an energy spread of 0.3 eV (**Figure 1b**). As a result, the SNRs at high spatial frequencies increase. Whereas the differences are small for resolutions worse than 2 Å, the use of the CFEG boosts the CTF from 58% to 91% at 1.5Å, and from 26% to 80% at 1.2Å. Since the SNR scales with the square of the CTF, this leads to a 2.5-fold increase in the SNR at 1.5 Å and a 9.5-fold increase at 1.2 Å (**Figure 1c**, **Methods**).

A disadvantage of operating the FEG at room temperature is that gas atoms can absorb to the surface of the emitter tip, which reduces the number of electrons generated. The contamination builds up over time and can be removed by periodically heating the tip. This procedure is called tip flashing and takes less than a minute. By performing tip flashing every 6-10 hours, based on automated monitoring of the beam current in the data acquisition program EPU, we found that the CFEG beam current remains stable over multiple days (**Extended Data Figure 1**).

### A stable energy filter

When electrons pass through the specimen, two types of interactions occur. Elastically scattered electrons maintain their original energy and contribute to the signal in the images. Inelastically scattered electrons deposit part of their energy in the sample, which, if left untreated^16^, contribute noise to the images. Therefore, SNRs can be improved by filtering away electrons that have experienced an energy loss. For this purpose, we used a new energy filter that TFS developed using the experience that previously led to the energy filter reported by Kahl et al^17^. This filter, located below the microscope’s projector column, comprises a 90° bending prism, an adjustable energy selecting slit and multiple electromagnetic lenses to correct for aberrations and to enlarge both the energy dispersion and the image. The design of the new filter was optimised for optical stability and reproducibility. The mechanics were designed to minimise the impact of temperature variations on the position of the optical elements, including the energy slit, with respect to the optical axis of the system. The prism was designed to have a large bending radius (135 mm), which reduces third and fourth order distortions. The new filter is stable over many days of operation: the position of the electrons that have not lost any energy, the so-called zero-loss peak, with respect to the centre of the energy slit changes less than 3 eV in either direction (**Figure 1d**).

### A next-generation camera

The Falcon series of direct-electron cameras are composed of 4096 × 4096 detector pixels, each measuring 14 × 14μm^18^. Their relatively large pixel size permits using a thick epilayer, which benefits the signal to noise ratio^19,20^. Since the third generation Falcon (Falcon-3), reset noise has been minimized by multi-frame correlative double sampling, so the noise in the resulting images arises mainly from the variation in the height and the shape of the charge distribution that is generated by individual incident electrons, or events. This noise may be reduced by a counting algorithm that positions a constant signal distribution at an estimated centre position for each event. But in order to identify all individual events, their proximity on each frame should be minimized. The Falcon-1 and Falcon-2 cameras, with a frame rate of 18 Hz, could only be operated in a charge-integrating mode. With a frame rate of 40 Hz, the Falcon-3 could also be operated in electron-counting mode, but only when spreading the electron dose over typical exposure times of close to a minute, thus restricting data collection throughput.

The latest Falcon-4 camera operates at 248 Hz, while an improved counting algorithm estimates event positions with sub-pixel accuracy and a better epilayer design optimises the event size versus event signal strength and thereby reduces the number of events that are lost during counting (**Figure 1e**). As a result, the Falcon-4 records images with typical exposure times of a few seconds, while the detective quantum efficiency is similar to the Falcon-3 in counting mode (**Figure 1f**). In addition, the resulting movies can be written out in a new data format, called electron-event representation (EER)^21^. Whereas conventional movie formats require a decision about dose fractionation into a limited number of movie frames at the time of data acquisition, the EER-format stores each detected electron event as a x,y-position on a four-times oversampled grid and its original time, or frame, of recording. Thereby, dose fractionation can be decided during image processing and modified depending on the task at hand. We adapted our image processing software RELION^22^ (see **Methods**) to be able to read the new data format, thus allowing motion correction by the Bayesian polishing algorithm^23^ and reconstructions of movie frames with the original 248 Hz time resolution of the camera.

### Application to a human GABA_A_ receptor sample

To evaluate the impact of the technological advances described above in a systematic manner and on a challenging sample, we focused on a ~200 kDa human membrane protein, a prototypical GABA_A_ receptor (GABA_A_R) β3 homopentameric construct bound to its small molecule agonist, histamine^3,24^. GABA_A_ receptors, the major inhibitory neurotransmitter receptors in the vertebrate brain, are a large family of pentameric ligand-gated chloride channels that carry the primary target sites for a wide range of clinically-relevant drugs including general anaesthetics, benzodiazepines, barbiturates and neuroactive steroids^25^. Despite their widespread use, many of these compounds exhibit unwanted side effects. Yet, structure-based drug discovery of more specific modulators has been hampered by the difficulty of solving high resolution GABA_A_R structures, with maps achieving 2.5-3 Å resolution only in the best-case scenarios^26^. While physiological GABA_A_Rs are typically heteromeric, the homomeric construct used here, bound to the megabody Mb25^24^, eliminates difficulties related to sample and grid preparation and its five-fold symmetry allowed us to collect smaller datasets and vary data collection strategies in a time-efficient manner.

We illustrate these tests with six data sets, collected from three different GABA_A_R grids on three different Titan Krios microscopes (**Extended Data Table 1**) and assessed their quality in terms of their estimated *B*-factors^10^ (**Methods**, **Figure 2a**). To assess the current state-of-the art, we collected a data set on a microscope with an XFEG and a bottom-mounted (BM) Falcon-3 camera (without energy filter), which resulted in a *B*-factor of 72 Å^2^. An improvement of 15 Å^2^ in the *B*-factor was observed when we imaged a second grid on a microscope with an XFEG, the new energy filter (with retracted slits) and a Falcon-4 camera. Using the same grid and the same microscope, a further improvement of 7 Å^2^ in the *B*-factor was achieved when using the energy filter with a slit width of 5 eV. Varying the energy slit widths to 3eV and 10 eV did not have a noticeable effect on the *B*-factors (**Extended Data Figure 2a**). Imaging a third grid on yet another microscope, with a CFEG, the new energy filter (with a slit width of 5eV) and a Falcon-4 camera, led to a small improvement for those reconstructions with resolutions beyond 2 Å. The latter is expected from the relatively small gains in the CTF envelope at those resolutions when using a CFEG instead of an XFEG (**Figure 1c**).

**Figure 2:**
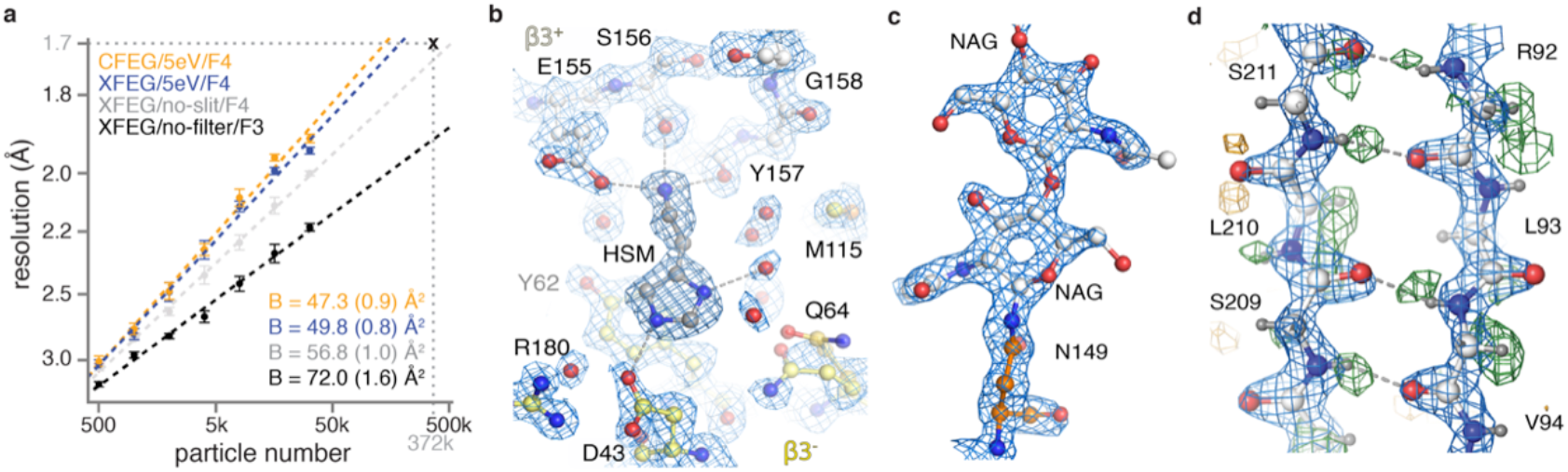
GABA_A_R reconstructions. **(a)** B-factor plots for four data sets using: the new CFEG, the new energy filter with a slit width of 5eV and a Falcon-4 camera (orange); an XFEG, the new energy filter with a slit width of 5eV and a Falcon-4 (blue); an XFEG, the new energy filter with the slits retracted and a Falcon-4 (grey); and an XFEG, no energy filter and a (bottom-mounted) Falcon-3 detector (black). B-factors estimated from the slope of fitted straight lines are shown in the same colours, with their standard deviations between brackets. **(b)** Overview of map quality at the agonist binding pocket, illustrating HSM coordination and a large number of water molecules (red spheres). **(c)** The N-acetyl glucosamine moieties attached to Asn149. **(d)** Difference map (green positive; orange negative) visualises hydrogen atoms in the hydrogen bonding network between β-strands.

By combining the energy-filtered data sets on the XFEG and the CFEG, we obtained a 1.7 Å resolution reconstruction from 371,693 particles (**Extended Data Figures 2** **and** **3**, **Extended Data Table 1**). The resulting map provides unprecedented levels of detail for a protein structure, other than apo-ferritin, solved by single-particle cryo-EM (**Extended Data Figure 3b-c**). The five symmetry-related agonist pockets, located at β3^+^-β3^−^ interfaces, are each occupied by one histamine (HSM) molecule with its imidazole ring stacked between the side chains of Phe200 and Tyr62 and its ethylamine chain nitrogen engaged in hydrogen bonds with the Glu155 carboxyl and backbone carbonyls of Ser156 and Tyr157, and a cation-π interaction with Tyr205 (**Figure 2b**, **Extended Data Figure 3d**). This binding mode is reminiscent of that observed for benzamidine^3^ and the neurotransmitter GABA^26^ (only in the nanodisc structures). However, for the first time, it is now possible to visualize the coordinating carboxyl group of Glu155, the carboxyl group of Asp43 coordinating an imidazole nitrogen of HSM, alongside a large network of ordered water molecules that surround the ligand to fill the agonist pocket (**Figure 2b**). Nevertheless, difference maps calculated in REFMAC^27^ (**Extended Data Figure 3e**) indicate that the density for the carboxyl groups of both Asp43 and Glu155 is too weak, suggesting that he pocket may be affected by radiation damage. The latter is confirmed by a movie of the dose-dependent evolution of the cryo-EM density in the ligand pocket (**Extended Data Movie 1**). Similarly, the disulfide bond between Cys136 and Cys150, a signature of the Cys-loop receptor family of which GABA_A_R is a member, appears to be partially reduced during the electron exposure (**Extended Data Figure 3f**, **Extended Data Movie 2**).

Noteworthy are also the exceptional quality of densities for N-linked glycans attached to Asn149 (**Figure 2c**), underlying their functional importance in providing structural support for the agonist-binding “loop C” and facilitating signal transduction between the extracellular and transmembrane domains of the receptor. A large number of amino acid side chains occupy alternative conformations while, in the most ordered regions of the receptor, hydrogen-bonding networks can be visualised directly in difference maps with a refined model from which the hydrogen atoms were omitted (**Figure 2d**, **Extended Data Figure 3d**). Multiple lipid molecules surround the transmembrane region of the receptor (**Extended Data Figure 3h-i**). Similar to recent observation for the ~1.9 Å resolution connexin maps^28^, the lipid densities are less resolved relative to the protein, presumably reflecting the multiple alternative conformations they adopt in the nanodisc environment. Nevertheless, key positions for GABA_A_R modulation including the general anaesthetics pocket29 and the site of neurosteroid modulation^30,31^ are clearly occupied by lipid tails (**Extended Data Figure 3j-k**), raising important questions for future mechanistic investigation.

### Atomic resolution structure of apo-ferritin

To explore the potential of the new technologies to yield even higher resolution reconstructions, we tested a mouse apo-ferritin sample. Apo-ferritin is a popular cryo-EM benchmark because its molecular stability and its 24-fold symmetry allow high-resolution reconstructions to be calculated from relatively few particles. We previously reported a reconstruction of human apo-ferritin with a resolution of 1.65 Å from 426,450 particles recorded on a Titan Krios microscope with a Gatan energy filter and a Gatan K2 camera^22^. The *B*-factor for that data set was 66 Å^2^.

Using the same microscope with a CFEG we employed for the GABA_A_ receptor, we imaged mouse apo-ferritin on the Falcon-4 camera, with a slit width of 10 eV in the new energy filter. From 3,370 movies, we obtained a reconstruction with a resolution of 1.2 Å from 363,126 particles (**Extended Data Figure 4**). Corrections during image processing, for third and fourth-order optical aberrations^32^, as well as for the curvature of the Ewald sphere^22,33^, were crucial in obtaining a *B*-factor of 32 Å^2^ (**Figure 3a**). After global sharpening of the map following standard procedures, most atoms are resolved as individual blobs of density; C=O bonds and some of the C-N bonds in the peptide planes are less well separated (**Figure 3b-d**). Many hydrogen atoms, including those from individual water molecules, are resolved in a REFMAC difference map, more clearly than in the GABA_A_ receptor case and allowing a direct analysis of hydrogen bonding networks (**Figure 3e-f**).

**Figure 3:**
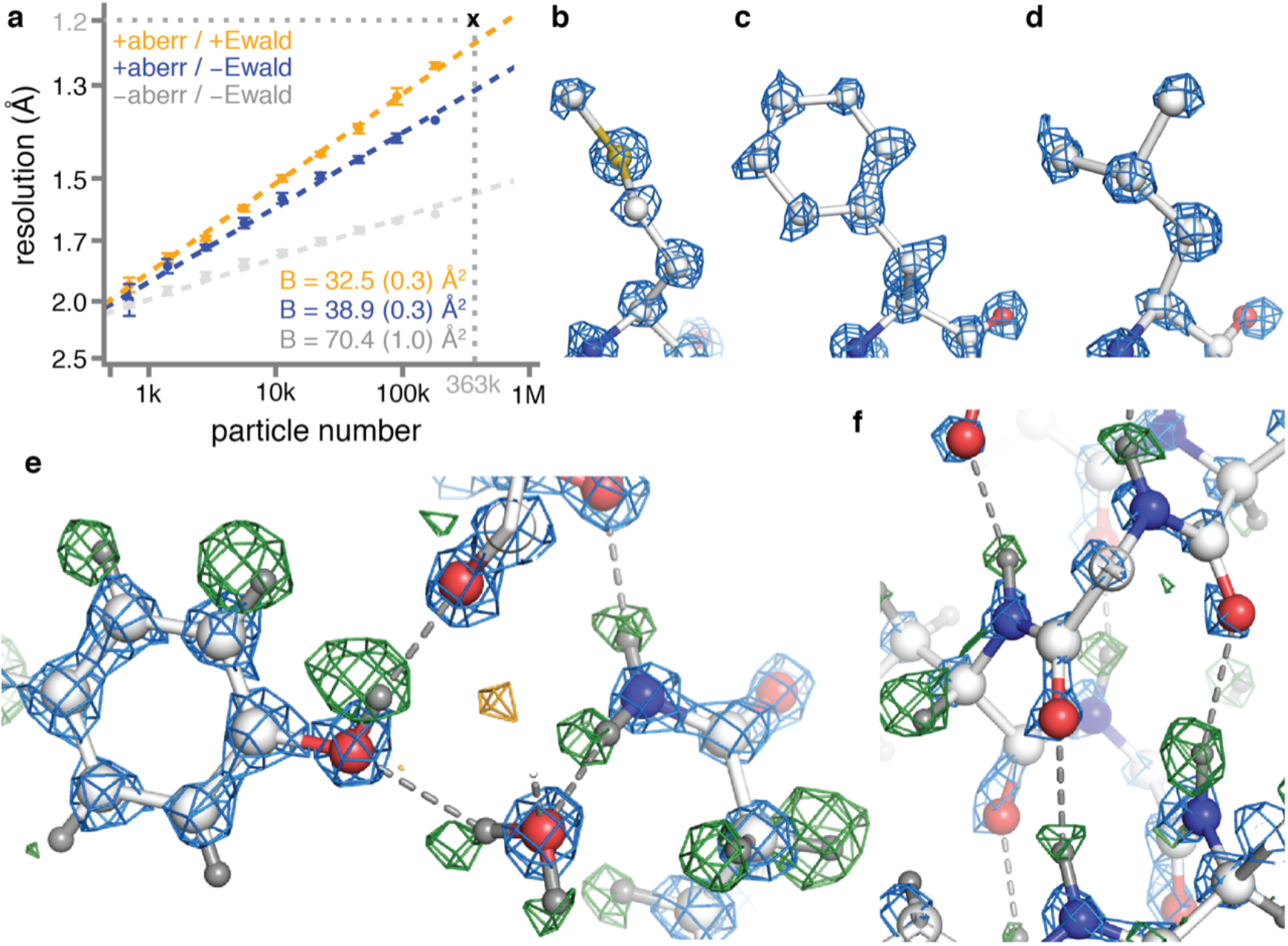
Apo-ferritin reconstruction. **(a)** B-factor plots for reconstructions using: high-order aberration and Ewald sphere correction (orange); high-order aberration correction only (blue); and no correction (grey). B-factors estimated from the slope of fitted straight lines are shown in the same colours, with their standard deviations between brackets. **(b)** Density for M100 in the 1.2 Å map is shown in blue. **(c)** Density for F51. **(d)** Density for L175. **(e)** Hydrogen bonding network around Y32 and water-302 is visible in difference map (green positive; orange negative) **(f)** As in (e), for hydrogen bonding network in the α-helix at residues ^21^NRQIN^25^.

We also refined an atomic model with hydrogen atoms using a so-called riding model, in which the lengths of chemical bonds involving hydrogens are constrained to expected values. When analysing the resulting hydrogen positions, we noticed a mismatch between the centre of the density in the difference map and the position of the hydrogen atoms in the model. The density peaks in the difference map were consistently further away from the bonded atom than the hydrogen atoms at its expected position. This result is explained by the observation that electron imaging visualises electrostatic potential, which depends both on the positions of nuclei and electrons. The magnitude by which the observed electrostatic potential density will be shifted away from the proton position will depend on the resolution of the map and the local order as modelled by atomic B-values (see **Methods**, **Extended Data Figure 5**). These observations suggest that cryo-EM reconstructions to atomic resolution may also be used to visualise different chemical bonding effects^34,35^, which is a subject for future investigations.

Finally, we used the EER movies to calculate reconstructions for each of the 434 individual camera frames that were acquired per exposure. Thereby, we were able to follow the dose-dependent evolution of the cryo-EM density with a resolution of 0.1 electrons (e^−^) per Å^2^ (**Extended Data Movie 3**). During the first ~0.3-0.4 e^−^/Å^2^ the reconstructions are of relatively poor quality, probably because our motion correction algorithm did not model the initial beam-induced motions adequately. As radiation damage is minimal in the earliest part of the exposure, these observations suggest that further improvements in beam-induced motion, possibly by considering particle rotations and height changes, may still yield considerable improvements in attainable resolution. Following the initial 1.5 e^−^/Å^2^, the resolutions of the individual frame reconstructions extend to 1.7 Å and remain better than 2 Å up to an accumulated dose of 11.5 e^−^/Å^2^. Thereby, this type of analysis provides unprecedented resolution, in spatial frequency and in dose, for future developments to model electron radiation damage events in proteins.

### Outlook

Since the introduction of direct-electron cameras in 2013, technologies underlying single-particle cryo-EM have continued to improve. Among these, electron microscopes have become more efficient^4,36^; cameras have increased in speed and sensitivity^37,38^; more stable sample supports became available^39,40^; and new image processing algorithms have been developed^9,22,41–43^. Currently, cryo-EM structures with resolutions in the range of 3-4 Å are common^44^. Resolutions in the range of 1.5-2 Å, however, have been restricted to standard benchmark proteins, such as apo-ferritin^22,45^ and β-galactosidase^46^, and other highly symmetric samples, including an adeno-associated virus serotype 2 (AAV2) icosahedral capsid variant^47^, the dodecameric urease from *Y. enterocolitica*^48^ and sheep lens connexin^28^.

The three developments in electron microscopy hardware described in this paper provide a step-change in the achievable resolution by single-particle cryo-EM. Using apo-ferritin, we show that these developments allow structure determination of a protein to true atomic resolution, as per the Sheldrick criterion^49,50^. The improved energy spread of the CFEG plays a crucial role in extending resolutions to better than 1.5 Å, which is in agreement with similar observations by others^15^. While a similar, or even better, effect could be achieved through the use of a monochromator^51^, the experimental setup would be considerably more complicated compared to the CFEG strategy described here. Besides the importance of the improved energy spread, we find that image processing corrections for higher-order optical aberrations and for the curvature of the Ewald sphere are also essential in reaching atomic resolution with reasonable amounts of data.

Our results on the GABA_A_R illustrate how the new technology can improve cryo-EM structures beyond the highly stable test samples that apo-ferritin represents. In particular, the new energy filter and the Falcon-4 camera also provide significant improvements of SNRs at lower resolutions. Although variations between microscopes and grids can complicate direct comparisons of *B*-factors, our results on GABA_A_R suggest that using the Falcon-4 camera in combination with the new filter compared to using a bottom-mounted Falcon-3 camera has a considerable effect on the image quality, which leads to a *B*-factor improvement of 15 Å^2^. Inserting the slits of the energy filter further improves the *B*-factor by 7 Å^2^. As our grids are mostly monolayers of GABA_A_R molecules, we expect the typical ice thickness to be in the range of 200-400 Å. It is unclear whether the removal of inelastically scattered electrons from such a thin sample alone would be enough to account for this improvement, or whether other effects may play a role too.

The experiments described here were not optimised in terms of speed, as our focus was on resolution and the prototype Falcon-4 version did not write movies out at the highest possible rate. We typically collected two images per hole and physically moved the stage for each hole, resulting in an average data acquisition rates of 95 movies per hour (including Dewar filling times and CFEG flashing). Nevertheless, this was sufficient to collect the 3,370 apo-ferritin movies that resulted in the 1.2 Å map within 36 hours, and GABA_A_R datasets allowing ~2 Å reconstructions in ~9 hours. Fringe free imaging (FFI) and aberration-free image shift (AFIS), which allow acquisition of multiple images per hole and multiple holes per stage position, would allow considerably higher throughputs. This would be particularly beneficial in situations where large numbers of small molecule binders need to be screened. Moreover, the quality of the GABA_A_R map described here, allowing an unprecedented visualization of water molecules in the agonist-binding pocket alongside HSM, a ligand of only 8 non-hydrogen atoms, demonstrates that cryo-EM fragment-based drug discovery and design projects are now feasible.

The technology described here increases the SNRs in cryo-EM images, which will improve the orientational and class assignments of individual particles. This will generally increase the achievable resolution for difficult samples, including membrane proteins in micelles or lipid bilayers, small protein particles, structurally heterogeneous macromolecular complexes, as well as cellular tomography.

## Acknowledgements

We thank Mike van Beers, Marcel Veerhoek, Roland Jonkers for maintaining the Titan Krios microscopes; Walter van Dijk, Bram van de Kerkhof, Stan Konings and Gijs van Duinen for advice on optics and microscope alignments; Bart van Knippenberg, Andreas Voigt and Fanis Grollios for support with EPU software; Adrian Koh, Toby Darling and Jake Grimmett for support with computing; Garbi Lezcano Singla, Erik Franken for support with the EER format; and Gerald van Hoften and Gerard Hosmar for support with the Falcon-4 camera. This work was supported by the EM facilities at the MRC-LMB, the Biochemistry Department of Cambridge University and Thermo Fisher Scientific. The Cryo-EM Facility at Department of Biochemistry is funded by the Wellcome Trust (206171/Z/17/Z; 202905/Z/16/Z) and the University of Cambridge. We acknowledge funding from the UK Medical Research Council (MC_UP_A025_1012 to G.M., MR/L009609/1 and MC_UP_1201/15 to A.R.A. and MC_UP_A025_1013 to S.H.W.S.); the Japan Society for the Promotion of Science (Overseas Research Fellowship to T.N.); MRC-LMB, Cambridge Trust and School of Clinical Medicine, University of Cambridge (LMB Cambridge Scholarship and Cambridge MB/PhD fellowship to A.S.); the European Commission (Marie Skłodowska-Curie Actions H2020-MSCA-IF-2015/709054 to L.M. and H2020-MSCA-IF-2017/793653 to P.M.G.E.B.); EMBO (long term fellowship 300-2015 to L.M.); Cancer Research UK (T.M., grants C20724/A14414 and C20724/A26752 to Christian Siebold, University of Oxford); Boehringer Ingelheim Fonds (PhD Fellowship to J.M.).

## Data availability

Atomic coordinates for the GABA_A_R construct and apo-ferritin will be deposited in the Protein Data Bank; the cryo-EM density maps in the Electron Microscopy Data Bank; and the raw movies in the Electron Microscopy Public Image Archive. Meanwhile, the models and maps, as well as the Extended Data Movies mentioned in the main text can be downloaded from: ftp://ftp.mrc-lmb.cam.ac.uk/pub/scheres/atomic/

## Author contributions

E.d.J., J.K., M.B., J.M., and P.T. contributed to microscope hardware and software developments; G.M. performed DQE analysis; D.K., A.S. and S.M. prepared cryo-EM grids, A.K., L.Y., E.V.P, S.W.H and D.Y.C. acquired cryo-EM data; T.N., A.K, A.S., L.Y., S.M., P.M.G.E.B., I.T.G., L.M., J.M. and S.H.W.S analysed cryo-EM data; T.N., A.R.A., T.M. and G.M. prepared atomic models; T.N., A.K., A.R.A. and S.H.W.S. conceived the project and designed the experiments; G.M., A.R.A. and S.H.W.S. wrote the manuscript, with input from all authors.

## Methods

### Characterisation of the new microscopy hardware

The energy spread of the field emission gun was measured using the new energy filter. Post-slit multi-poles of the filter were set to spectroscopy mode in order to form an enlarged image of the spectrum on the camera. The intensity of the incoming beam was carefully reduced in order not to saturate or damage the camera. The energy dispersion at the camera was 8 meV/pixel. The exposure time was 31 ms. The energy spread at half the maximum value (FWHM=0.71 for XFEG and FWHM=0.28 for CFEG, **Figure 1b**) were used to calculate CTF envelope function, Δ*f*, using the following equation^51^:

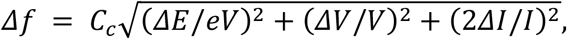

using *C*_*c*_=2.7mm; Δ*V/V*=0.02ppm and Δ*I/I*=0.1ppm. Note that the latter two contributions account for less than 1% drop in the CTF envelope.

The stability of the energy filter was assessed by repeatedly measuring the position of the zero-loss peak with respect to the energy-selecting slit. To this end, we mechanically moved one slit edge to the optical axis and fine-tuned an offset on the high tension supply such that the slit edge blocked half of the beam intensity. The resulting offset of the high tension was read out as the shift in the zero loss position. Measurements, each taking approximately 40 seconds, were repeated continuously for 14 days.

Dose response curves for the Falcon-3 and Falcon-4 cameras were measured using the flu-screen and fixed magnification steps. Before each experiment, the flu-screen was carefully calibrated with a Faraday cup in its linear range (0.4-5.0 nA at 300 kV). The read-out from the flu-screen was interpreted as the expected electron dose and plotted against the counted number of electrons on the camera.

DQE measurements were performed as described^52^ at 300 keV on a Titan Krios G2 microscope at the LMB for the Falcon-3 and on a Titan Krios G3i microscope at the Materials Science Department of the University of Cambridge for the Falcon-4. For calibration of the flu-screen on the microscope with the Falcon-4, a Faraday cage and a Keithley 6485 picoammeter (kindly provided by Gerard van Hoften, Thermo Fisher Scientific) were used. For the microscope with the Falcon-3, the drift tube of a Gatan energy filter (GIF; BioQuantum 686) was used as a Faraday cup and the current was measured using a Deben 087-001 SEM Probe current meter. Because the beam currents used in counting mode imaging are too low to be accurately measured, a defined beam with a measurable current was set at a low magnification and the desired electron flux on the camera was obtained by increasing the magnification. The relative magnification change was measured by using a higher current to fully illuminate the flu-screen at the low magnification and noting the change in current upon going to the higher magnification.

To account for the non-linearity that is introduced by coincidence loss, DQEs were first measured at a low dose rate (0.19 e^−^/pixel/s for the Falcon-3; 0.32 e^−^ /pixel/s for the Falcon-4) and then extrapolated to a typical dose rate (0.5 e^−^ /pixel/s for the Falcon-3; 3.6 e^−^/pixel/s for the Falcon-4) using the measured response versus dose-rate curves (**Figure 1e**). Modulation transfer functions (MTFs) were measured using the knife-edge method from a straight part of a shadow of the pointer on the camera. The actual area of the images used was chosen carefully by looking at the difference between the observed image and that predicted from the inferred MTF, to ensure that the edge in the selected area was straight and free of defects. In the case of the Falcon-4, the only part of the pointer edge that was suitable was at a 15 degrees angle with the pixel array of the camera, the effect of which was accounted for when calculating the MTF. Even at the lowest dose rates used, there was still a small decrease in the power spectra at low-spatial frequency due to coincidence loss. To avoid overestimating the resulting DQE at low spatial frequency, the noise power spectra were extrapolated from a Gaussian fit of the power between 0.2 and 0.8 of Nyquist. The amplitudes and widths of fitted Gaussians to the noise power spectra were observed to be in excellent agreement with those expected from the number of counted electrons per frame and estimates from the point spread function of images of individual incident electrons.

### GABA_A_ receptor production, purification and nanodisc reconstitution

Human GABA_A_ receptor β_3_-subunit with a truncated M3-M4 loop was expressed in HEK293S cells^3^. Frozen cell pellets were resuspended in 50 mM HEPES pH 7.6, 300 mM NaCl, 1 mM histamine supplemented with 1% (v/v) mammalian protease inhibitor cocktail (Sigma-Aldrich) and solubilised with 1% (w/v) lauryl maltose neopentyl glycol (LMNG) and 0.1% (w/v) cholesteryl hemisuccinate tris salt (CHS). (No CHS was added to the sample that was used for the CFEG dataset). Receptors were purified with streptavidin-binding peptide (SBP) affinity chromatography. After the binding step, while the receptor was still bound to the beads, an excess of bovine brain extract (BBE; type I, Folch fraction I, Sigma-Aldrich, 20 mg/mL stock) and phosphatidylcholine (POPC, Avanti, 10 mg/mL) mixture (15:85, v/v) was added to the sample and incubated for 30 min then washed. Receptors were then reconstituted into membrane scaffold protein 2N2 (MSP2N2) lipid nanodiscs by incubating with an excess of MSP2N2 (0.6 mg/mL final concentration) and bio-beads for 2 h. Resin was then washed extensively and eluted with 2.5 mM biotin, 12.5 mM 4-(2-hydroxyethyl)-1-piperazineethanesulfonic acid (HEPES) buffer at pH 7.6, with 75 mM NaCl and 1 mM histamine.

### Cryo-EM grid preparation

Purified GABA_A_R was incubated with ~1 μM Megabody-25^24^ and 3.5 μl sample was applied to glow discharged 300 mesh 1.2/1.3 and 2/2 UltraAuFoil gold grids (Quantifoil) for 30 s then blotted for 5.5 s prior to plunge-freezing with liquid ethane cooled by liquid nitrogen. Plunge-freezing was performed using a Leica plunger (Leica Microsystems; XFEG datasets) or Vitrobot Mark IV (Thermo Fisher Scientific; CFEG datasets) at 100% humidity and 14 °C.

A frozen aliquot of 7mg/ml mouse apo-ferritin in 20mM HEPES pH 7.5, 150mM NaCl, 1mM dithiothreitol (DTT) and 5% trehalose, which we received from the Kikawa Lab at Tokyo University, was thawed at room temperature and cleared by centrifugation at 10,000g for 10 min. The supernatant was diluted to 5mg/ml with 20mM HEPES pH 7.5 150mM NaCl, and 3 μl of the diluted sample was applied onto glow-discharged R1.2/1.3 300 mesh UltrAuFoil gold grids (Quantifoil) for 30 s and then blotted for 5 s before plunge-freezing the grids into liquid ethane cooled by liquid nitrogen. Plunge-freezing was performed using a Vitrobot Mark IV (Thermo Fisher Scientific) at 100% humidity and 4 °C.

### Cryo-EM data acquisition

All cryo-EM data were collected on Falcon cameras in electron counting mode using Titan Krios microscopes (Thermo Fisher Scientific) operating at 300 kV. Before data acquisition, two-fold astigmatism was corrected and beam tilt was adjusted to the coma-free axis using the autoCTF program. All datasets were acquired automatically using EPU software (Thermo Fisher Scientific). Detailed data acquisition parameters for all data sets are given in **Extended Data Table 1**.

For GABA_A_R, one data set was acquired on a Titan Krios at the Department of Biochemistry, University of Cambridge, UK. This microscope is equipped with an XFEG and a bottom-mounted (BM) Falcon-3 camera. All other data sets were acquired at the Thermo Fisher Scientific RnD division in Eindhoven, The Netherlands. Four GABA_A_R data sets were collected on a Titan Krios equipped with an XFEG and a prototype of the Falcon-4 that was mounted behind the new energy filter. These datasets were collected consecutively over a period of 5 days from the same grid to study the effect of varying energy slit widths. To keep the ice thickness similar between these experiments, all the exposure areas were selected at the start using a constant ice filter within EPU software. One GABA_A_R data set was acquired on a microscope with a CFEG and a prototype of the Falcon-4 that was mounted behind the new energy filter. The slit width of the energy filter was set to 5 eV. For this data set, 5573 movies were acquired using a 100um objective aperture and 3160 movies were collected without an objective aperture. The CFEG was automatically flashed every 5 hours using the EPU software.

For the apo-ferritin data set, we used the same microscope with the CFEG, the new filter and the prototype of the Falcon-4 camera. No objective aperture was used and the slit width of the energy filter was set to 10 eV.

### The EER movie format

Electron event representation (EER) is a movie format that is introduced with the Falcon-4 camera^21^. Whereas in the existing MRC image format groups of subsequent camera frames, or dose fractions, are summed and represented as images, EER stores the location and time of recording of each detected electron event. The EER format records the event coordinates in a four times oversampled, that is a 16k×16k, grid. The time resolution is the original frame rate of the camera, so that dose fractionation during data acquisition is no longer needed. Instead, beam-induced sample motions or stage drift can be corrected without a loss in temporal resolution. The electron event stream is stored on disk after compression with a run-length encoding algorithm. For GABA_A_R, data sets were acquired at a dose rate of 4.7 electrons per pixel per second, at a pixel size of 0.727 Å and a total dose of 18.1 electrons per pixel. These settings yielded EER movies consisting of 1113 frames and with an average size of 485 MB. For apo-ferritin, data were acquired at a dose rate of 4.5 electrons per pixel per second, at a pixel size of 0.457 Å and a total dose of 40 electrons per Å^2^, resulting in movies of 434 frames occupying on average 160 MB.

To read EER movies, we modified the publicly available RELION-3.1-beta. Inside RELION, the events from the 16k×16k grid are positioned as binary pixels in a 8k×8k grid, which is then Fourier cropped into a 4k×4k grid. Using the modified version, EER movies can be used for frame alignment using RELION’s implementation of the MotionCor2 algorithm^53^ and for per-particle beam-induced motion correction using its Bayesian polishing program^23^. For both programs, the user needs to specify a dose fractionation rate.

Both the implementation of reading EER movies into RELION and the EER format itself are distributed as free, open-source software.

### Apo-ferritin image processing

A total of 3370 movies in EER format were motion corrected with RELION’s implementation of the MotionCor2 algorithm^53^. For this purpose, original hardware movie frames were dose fractionated into groups of 14 frames, corresponding to an accumulated dose of 1.3 e^−^/Å^2^ per fraction. CTF estimation was performed with CTFFIND-4.1.13^54^ using the sums of power spectra from combined fractions corresponding to an accumulated dose of 4 e^−^/Å^2^. Micrographs whose estimated resolution from CTFFIND was worse than 5 Å were removed, leaving 3080 micrographs for further processing.

Using the Laplacian-of-Gaussian algorithm, 428,590 particles were picked and subjected to 2D classification. After selection of good classes, 380,236 particles were subjected to standard 3D auto-refinement to give an initial reconstruction with an estimated resolution of 2.3 Å. Subsequently, three runs of CTF refinement^32^ were performed: first refining magnification anisotropy; then refining optical aberrations (up to the 4th order); and finally refining per-particle defocus and per-micrograph astigmatism. A 3D auto-refinement with the refined values yielded an estimated resolution of 1.7 Å. Bayesian polishing to optimise per-particle beam-induced motion tracks, followed by another round of auto-refinement resulted in a 1.43 Å map. CTF refinement was then repeated for optical aberration correction, magnification anisotropy, per-particle defocus and per-micrograph astigmatism. To compensate for changes in the tilt of the electron beam during the data acquisition session, separate optics groups were created for particles belonging to consecutive groups of 125 micrographs, or whenever the CFEG was flashed. At this point, 201 micrographs with refined scale factors (*rlnGroupScaleCorrection*) with values below 0.5 were also removed, resulting in 363,126 particles in the final set. Following a last round of auto-refinement, these particles were subjected to a second round of Bayesian polishing with a large particle extraction box (spanning 320 Å), such that high-resolution signals that are delocalised by the CTF were still contained in the images. At this point, seven hardware frames were grouped into each dose fraction to be able to capture more rapid motions. The resulting particles were then used for reconstruction with Ewald sphere correction^33^. The estimated resolution of the final map, following standard post-processing procedures in RELION, was 1.22 Å.

### GABA_A_ receptor image processing

GABA_A_R data sets were processed following the same strategy as for apo-ferritin, with the following exceptions. All movies, except for the ones acquired on the microscope with a CFEG, were recorded in MRC format. For the cold-FEG data, hardware movie frames were initially dose fractionated into groups of 24 frames, corresponding to an accumulated dose of 0.86 e^−^/Å^2^ per fraction. For the second round of Bayesian polishing, a dose fractionation of 12 hardware frames was used. Particle picking was performed using an in-house generated Linux port of the of BoxNet2D neural network in Warp^41^. Prior to the last auto-refinement, 3D classification into three classes without alignment was performed to select the particles contributing to the best reconstruction. The first part of the CFEG data was acquired with a 100 μm objective aperture. Charging of the aperture probably led to third and fourth order optical aberrations, which were estimated during CTF refinement in optics groups of 160 micrographs, or whenever the CFEG was flashed or liquid nitrogen was refilled. The second part of the CFEG data was acquired without an objective aperture. For these images, no significant higher-order optical aberrations were observed and creating multiple optics groups was deemed unnecessary. The mask used for resolution estimation during post-processing only contained density for GABA_A_R; all density corresponding to the mega-bodies was removed.

### B-factor estimation

For each GABA_A_R dataset, a random subset of 32,000 particles was selected from the final set of particles after the second round of Bayesian polishing. This subset was subjected to 3D auto-refinement and the resulting orientations were used to calculate reconstructions for each of the two random halves used in the auto-refinement. Similar half-set reconstructions were also calculated for random subsets of 16,000, 8,000, 4,000, 2,000, 1,000 and 500 particles. This procedure was repeated seven times. For each subset and repeat, resolution estimation was again performed using standard post-processing functionality in RELION. The square of the resulting estimated resolutions were then plotted against the natural logarithm of the number of particles in the subset, and B-factors were calculated from the slope of a straight line fitted through all points in the plot.

For apo-ferritin, the same procedure was used, with the exception that the orientations from the final refinement were used, and subsets were generated by halving the number of particles eight times, starting from a subset of 320,000 particles.

### Atomic modelling

The atomic models for GABA_A_R and apo-ferritin were derived from PDB-4cof and PDB-6s61, respectively. Both models were manually adjusted in COOT^55^. Difference maps were used to rebuild the model, add alternative conformations and waters. Real-space refinement was performed using PHENIX, version 1.16.354956 for GABA_A_R. Both models were refined with anisotropic atomic B-factors in reciprocal space using REFMAC, version 5.8.0258^27^. For cross-validation, the atomic models refined against the full map was perturbed by random shifts of the atoms of up to 0.3 Å and the perturbed model was refined in REFMAC against one of the two half-maps. FSC curves for the resulting model against that half-map (FSC_work_) and against the other half-map (FSC_free_) are shown with blue and dashed orange lines, respectively, in **Extended Data Figures 2** and **4**. Molprobity score, clash score, rotamer and Ramachandran analyses were performed using MOLPROBITY^57^.

### Hydrogen position calculations

X-ray scattering and electron scattering factors are related through the Mott-Bethe formula, which is the solution of Poisson equations in Fourier space under certain conditions:

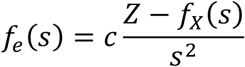

Where 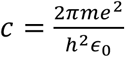; *m* is the mass of an electron; *e* is the charge of an electron, *h* is the Planck constant; and *ϵ*_0_ is the dielectric constant of the vacuum; *Z* is the nuclear charge; *f*_*X*_(*s*) is the X-ray scattering factor; *f*_*e*_(*s*) is the electron scattering factor; and *s*=*1/d* is the resolution. Adding the position-independent, isotropic atomic *B*-factor gives:

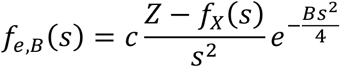

This formula is valid if the centre of electron density coincides with the position of the nucleus. If this is not the case, the formula should be modified according to the vector Δ*x* between the centre of electron density and the nuclear position (assuming that the centre of electron density is at the origin):

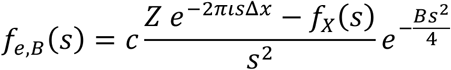

Note that, in general, the form factor for electron scattering becomes a complex quantity. For all non-hydrogen atoms, it is safe to assume that the centre of electron density and the nuclear position coincide.

For hydrogen atoms, the centre of electron density and the nuclear position do not coincide and we are interested in the peak height for the density of electrostatic potential along the bond between a hydrogen and its parent atom. The below ignores the fact that electron densities will be non-spherical, as they are elongated along the bond, thus adding additional complexity to Mott-Bethe formula.

The densities for the proton and the electron of a hydrogen atom at a resolution *s*_*max*_ and with a given *B* value is given by:

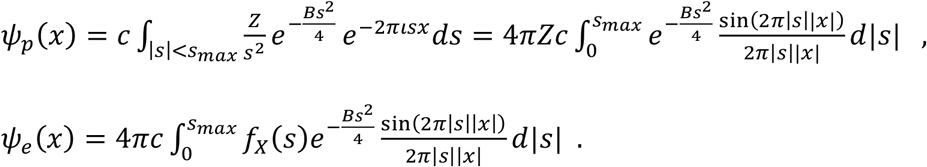

To calculate the density at a position *x* we need to calculate:

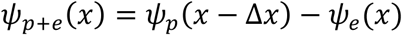

Even if both densities corresponding to the proton and the electron are spherically symmetric, the combined density will not be spherical. **Extended Data Figure 5** shows density profiles and their maxima for a hydrogen bonded to a carbon atom for different resolutions and B values. These profiles were calculated with the assumption that the electron-carbon distance is 0.98 Å and the proton-carbon distance is 1.09 Å.

---

**Extended Data Figure 1:**
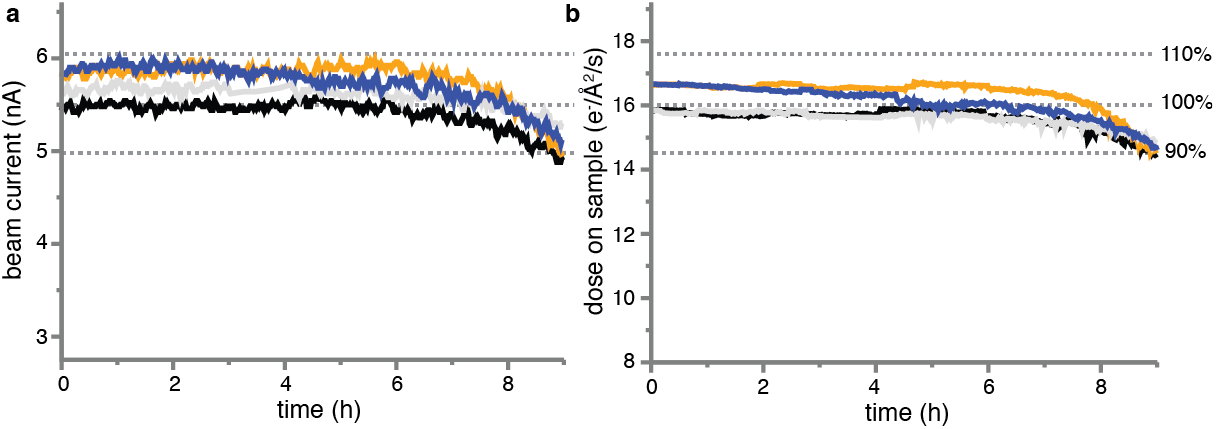
Characteristics of the new cryo-EM technology. **(a)** Four consecutive measurements of the CFEG beam current over a period of nine hours. The FEG tip was flashed just before the start of each measurement. **(b)** The dose at the sample as measured for the same four experiments in (a).

**Extended Data Figure 2:**
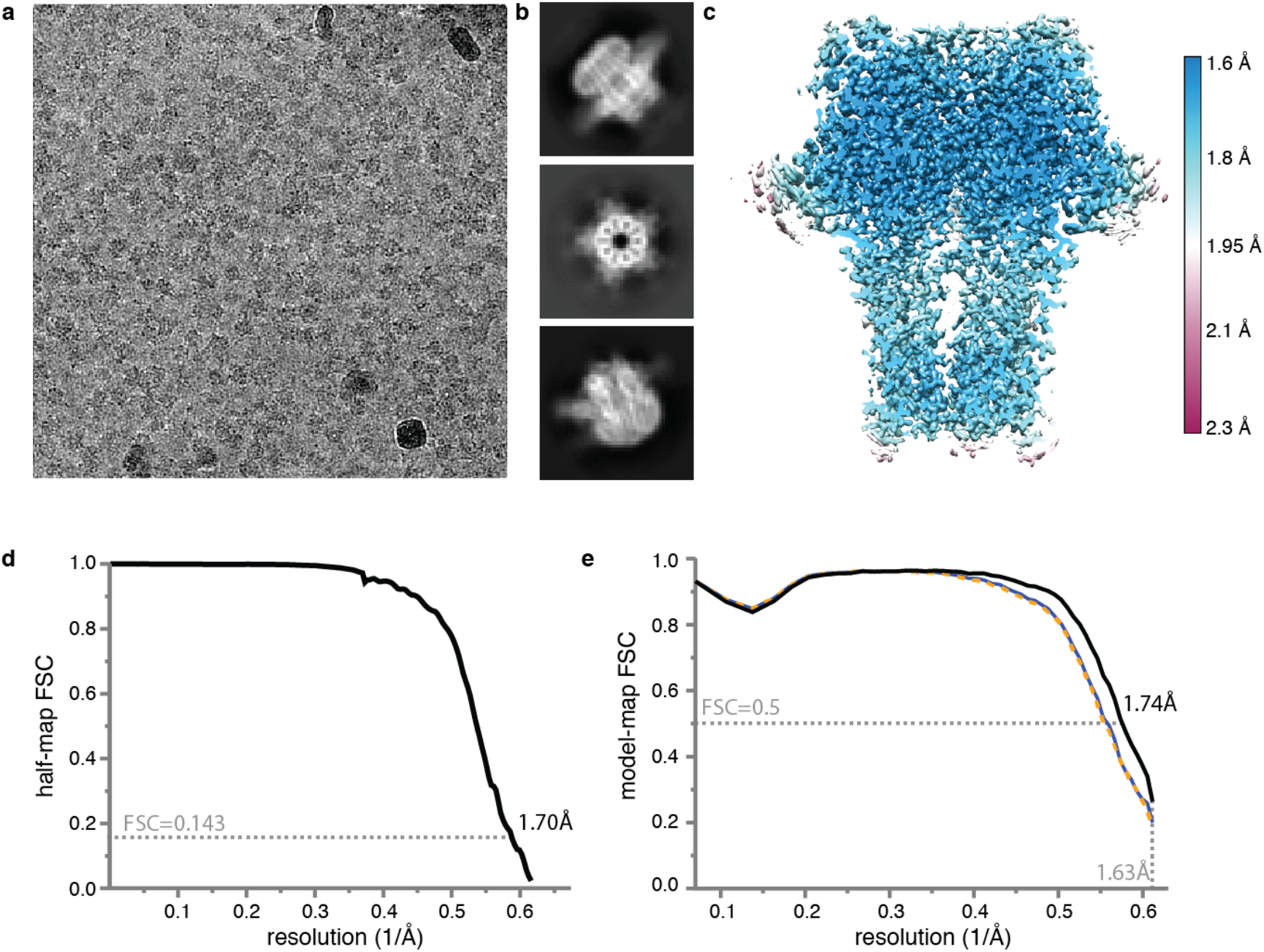
Cryo-EM for GABA_A_R. **(a)** Representative electron micrograph. **(b)** 2D class average images. **(c)** Local resolution map. **(d)** Fourier Shell Correlation (FSC) between the two independently refined half-maps. **(e)** FSC between the model and the map as calculated for the model refined against the full reconstruction against that map (black); the model refined in the first half-map against that half-map (FSC_work_; blue); and the model refined in the first half-map against the second half-map (FSC_test_; dashed orange). Atomic models were refined including spatial frequencies up to 1.63 Å.

**Extended Data Figure 3:**
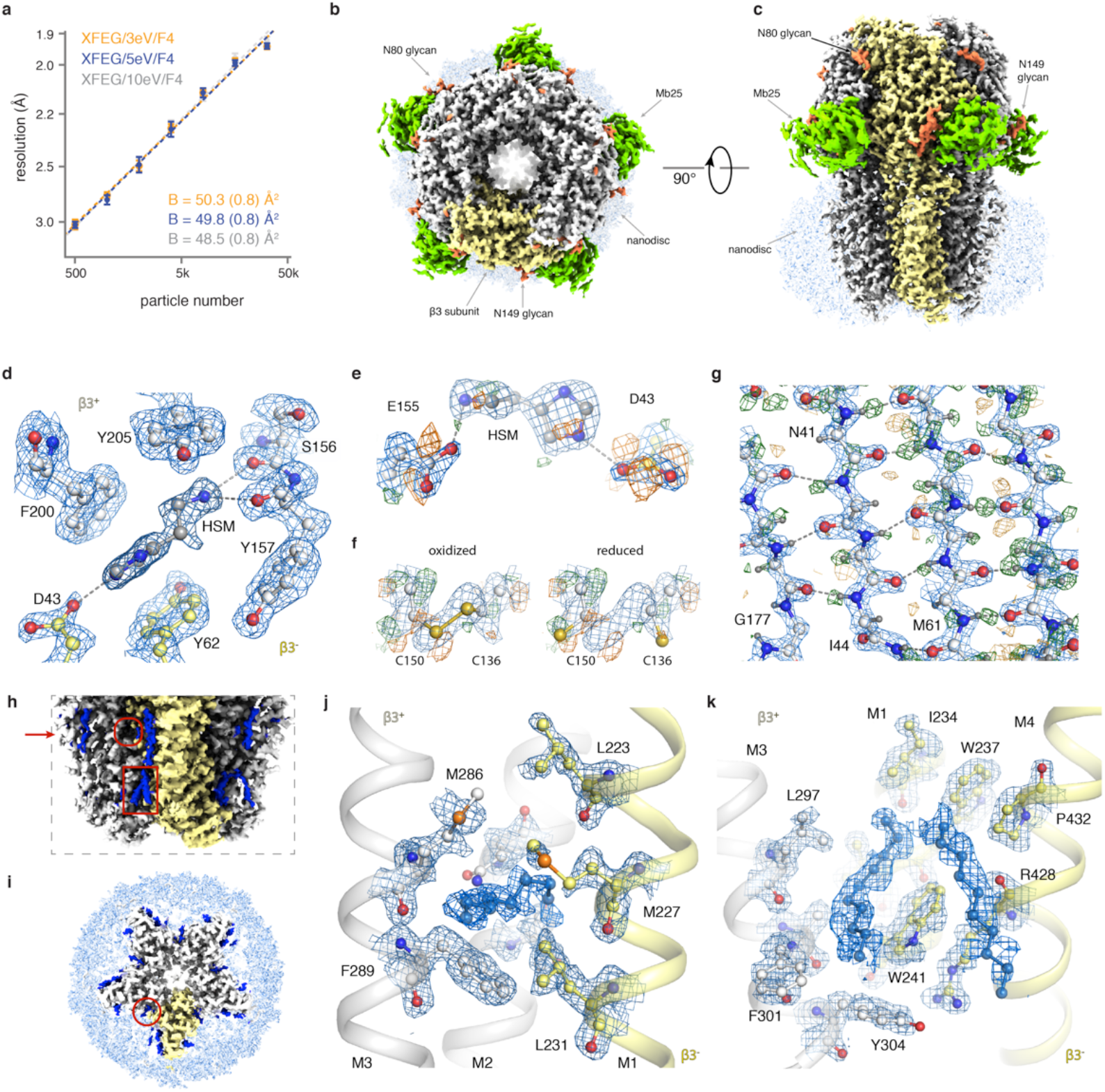
GABAAR reconstruction details. (**a**) B-factor plots for three data sets using an X-FEG, the new energy filter with a slit width of 3eV (orange), 5eV (blue) and 10 eV (grey) and a Falcon 4 camera. (**b**) The GABA_A_R cryo-EM map viewed from the extracellular space (top view). The density of one subunit of the homopentamer is highlighted in yellow; glycans are orange; the nanobody domain of Mb25 green; and the lipid nanodisc light blue. (**c**) As in b), but viewed parallel to the plasma membrane (side view). (**d**) Stacking of histamine with aromatic residues in the ligand-binding pocket. Water molecules removed for clarity. (**e**) Radiation damage in the ligand-binding pocket illustrated by difference maps of Asp43 and Glu155 carboxyl groups. (**f**) Radiation damage causes partial reduction of the disulfide bond between Cys136 and Cys150. Left panel: oxidized state, right panel: reduced state. Only Cα, Cβ and sulphur atoms depicted for clarity. (**g**) Difference map revealing the hydrogen bonding network between β-strands. (**h**) A close-up view of the lipids (blue) surrounding the transmembrane region of the receptor. Compared to (b-c), the contour level is decreased. Red arrow indicates section level as depicted in panel (i). The general anaesthetics pocket and the neurosteroid modulation site are indicated by the red circle and rectangle, respectively. (**i**) Top-view of a transverse section through the GABA_A_R transmembrane region shows semi-ordered lipids surrounding individual subunits. (**j**) Close-up view of the general anaesthetics pocket. (**k**) Close-up view of the neurosteroid modulation site. Lipids in (j) and (k) could not be identified unambiguously and therefore are modelled as aliphatic chains.

**Extended Data Figure 4:**
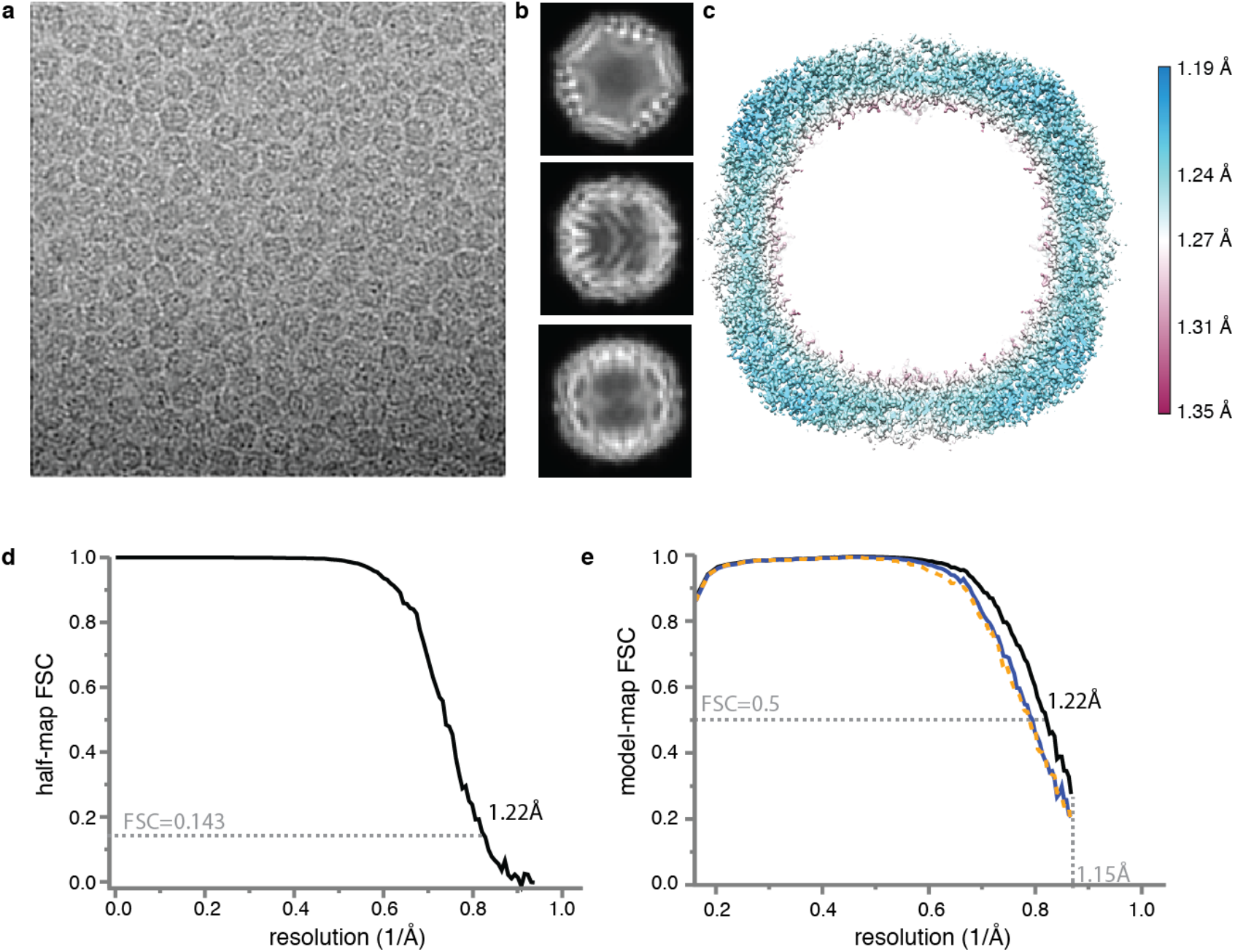
Cryo-EM for apo-ferritin. **(a)** Representative electron micrograph. **(b)** 2D class average images. **(c)** Local resolution map. **(d)** Fourier Shell Correlation (FSC) between the two independently refined half-maps. **(e)** FSC between the model and the map as calculated for the model refined against the full reconstruction against that map (black); the model refined in the first half-map against that half-map (FSC_work_; blue); and the model refined in the first half-map against the second half-map (FSC_test_; dashed orange). Atomic models were refined including spatial frequencies up to 1.15 Å.

**Extended Data Figure 5.**
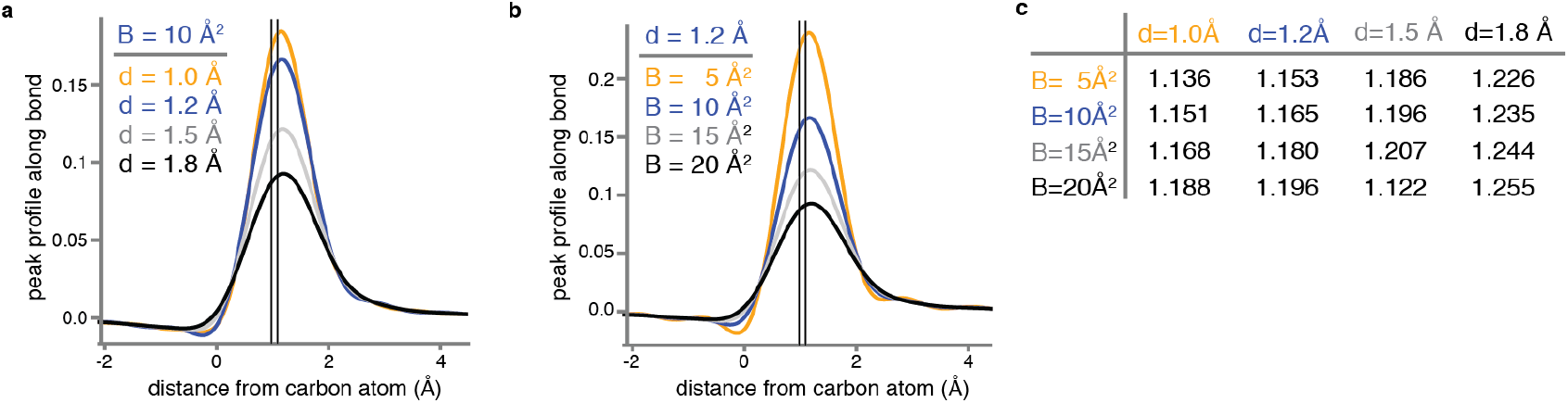
Electrostatic potential of hydrogen atoms. **(a)** Calculated profile (see Methods) of the electrostatic scattering potential along the bond between a carbon and a hydrogen atom, for a *B*-value of 10 Å2 and resolutions (*d*) of 1.0 Å (orange), 1.2 Å (blue), 1.5 Å (grey) and 1.8 Å (black). The two vertical lines indicate the electron-carbon distance of 0.98 Å and the proton-carbon distance of 1.09 Å. **(b)** As in (a), but for a resolution of 1.2 Å and *B*-values of 5 Å^2^, 10 Å^2^, 15 Å^2^ and 20 Å^2^. **(c)** Distances from the carbon atom (in Å) at which the calculated profile of the electrostatic scattering potential along the bond between a carbon and a hydrogen atom is at its maximum, for different resolutions (*d*) and *B*-values.

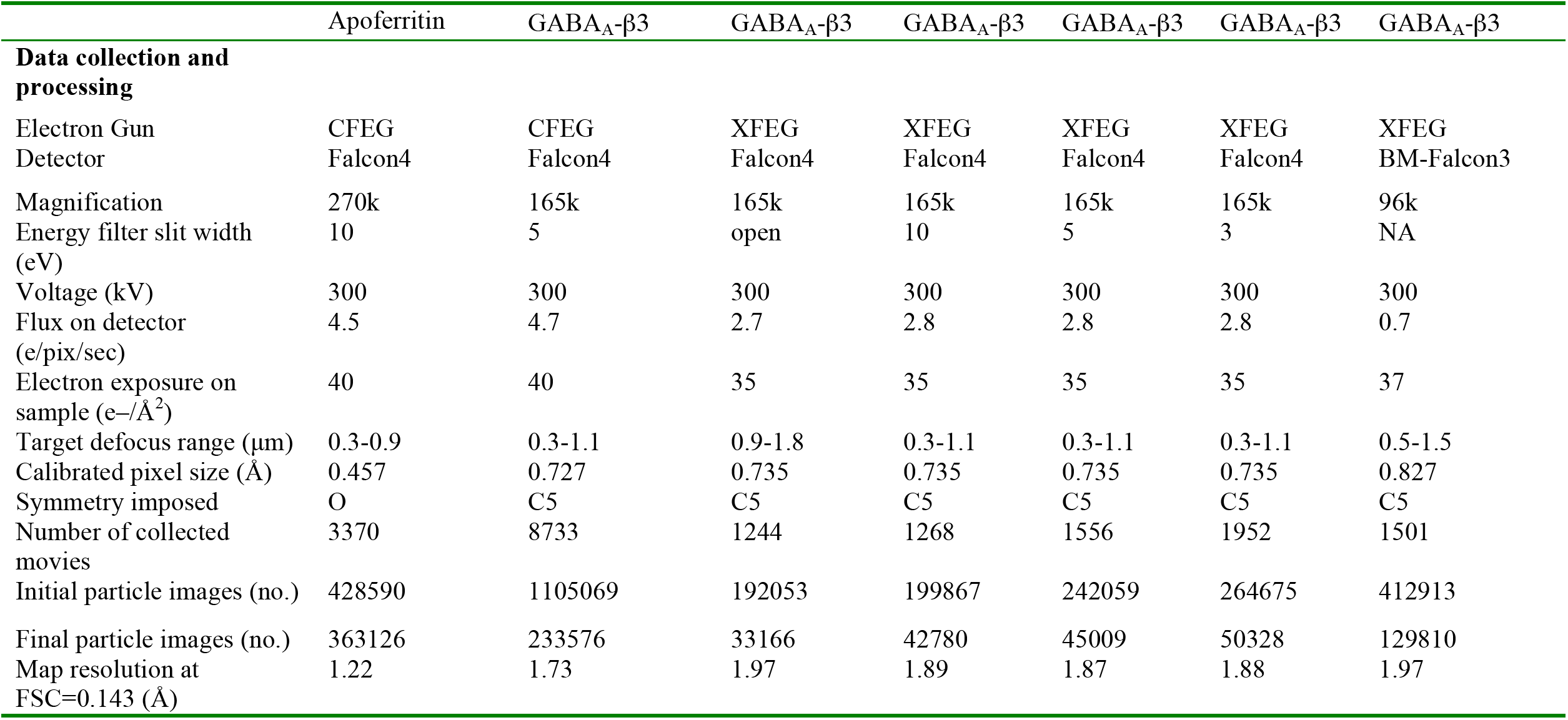
Cryo-EM data collection parameters

Cryo-EM data collection parameters
|  | Apoferritin | GABA<sub>A</sub>-β3 | GABA<sub>A</sub>-β3 | GABA<sub>A</sub>-β3 | GABA<sub>A</sub>-β3 | GABA<sub>A</sub>-β3 | GABA<sub>A</sub>-β3 |
| --- | --- | --- | --- | --- | --- | --- | --- |
| **Data collection and processing** |  |  |  |  |  |  |  |
| Electron Gun | CFEG | CFEG | XFEG | XFEG | XFEG | XFEG | XFEG |
| Detector | Falcon4 | Falcon4 | Falcon4 | Falcon4 | Falcon4 | Falcon4 | BM-Falcon3 |
| Magnification | 270k | 165k | 165k | 165k | 165k | 165k | 96k |
| Energy filter slit width (eV) | 10 | 5 | open | 10 | 5 | 3 | NA |
| Voltage (kV) | 300 | 300 | 300 | 300 | 300 | 300 | 300 |
| Flux on detector (e/pix/sec) | 4.5 | 4.7 | 2.7 | 2.8 | 2.8 | 2.8 | 0.7 |
| Electron exposure on sample (e-/Å<sup>2</sup>) | 40 | 40 | 35 | 35 | 35 | 35 | 37 |
| Target defocus range (μm) | 0.3-0.9 | 0.3-1.1 | 0.9-1.8 | 0.3-1.1 | 0.3-1.1 | 0.3-1.1 | 0.5-1.5 |
| Calibrated pixel size (Å) | 0.457 | 0.727 | 0.735 | 0.735 | 0.735 | 0.735 | 0.827 |
| Symmetry imposed | O | C5 | C5 | C5 | C5 | C5 | C5 |
| Number of collected movies | 3370 | 8733 | 1244 | 1268 | 1556 | 1952 | 1501 |
| Initial particle images (no.) | 428590 | 1105069 | 192053 | 199867 | 242059 | 264675 | 412913 |
| Final particle images (no.) | 363126 | 233576 | 33166 | 42780 | 45009 | 50328 | 129810 |
| Map resolution at FSC=0.143 (Å) | 1.22 | 1.73 | 1.97 | 1.89 | 1.87 | 1.88 | 1.97 |

Refinement and validation statistics

Refinement and validation statistics
| Refinement | Apoferritin<br>(EMDB-xxxx)<br>(PDB xxxx) | GABA<sub>A</sub>-β3<br>(EMDB-yyyy)<br>(PDB yyyy) |
| --- | --- | --- |
| Map resolution range<br>refined against (Å) | 1.22 - 136.5 | 1.6 - 291.1 |
| Map sharpening *B* factor<br>(Å<sup>2</sup>) | 2.3 | 23 |
| Model composition |  |  |
| Non-hydrogen atoms | 40512 | 21144 |
| Protein residues | 37584 | 19239 |
| Waters | 2880 | 1020 |
| Ions | 48 | 10 |
| Glycans | - | 515 |
| HSM | - | 40 |
| Lipids | - | 320 |
| *B* factors (Å<sup>2</sup>) |  |  |
| Protein | 25.3 | 46 |
| Waters | 39 | 40.8 |
| Ions | 33.4 | 50.5 |
| Glycans |  | 73.2 |
| HSM | - | 23.2 |
| Lipids | - | 105.4 |
| R.m.s. deviations |  |  |
| Bond lengths (Å) | 0.0155 | 0.009 |
| Bond angles (°) | 1.739 | 1.285 |
| Validation |  |  |
| MolProbity score | 1.69 | 1.17 |
| Clashscore | 4.96 | 2.69 |
| Poor rotamers (%) | 2.94 | 0 |
| Ramachandran plot |  |  |
| Favored (%) | 97.65 | 97.33 |
| Allowed (%) | 2.35 | 2.67 |
| Disallowed (%) | 0.00 | 0.00 |

RMS deviations are from REFMAC. Average *B* factors for GABA<sub>A</sub> calculated with average\_b.py plugin in PyMOL. Other values calculated by MolProbity from the PHENIX 1.17.1 package.

